# Novel reactivity phenotype mediates long-term consequences of cocaine exposure during adolescence and adulthood in male Sprague-Dawley rats

**DOI:** 10.1101/2025.07.10.664140

**Authors:** Antoniette M. Maldonado-Devincci, Rosalynn A. Amante, Ricardo Raudales, Laura A. Michael, Kimberly A. Badanich, Cheryl L. Kirstein

## Abstract

Individual differences in novelty seeking behavior often predict locomotor responses to psychomotor stimulants, an effect that may have a lasting impact throughout adolescence. The present study examined the role of novel reactivity phenotype on locomotor responses to repeated cocaine exposure during adolescence and adulthood in male rats. Prior to drug exposure, reactivity to a novel object was measured and animals were separated into low responders (LR) and high responders (HR). Cocaine was administered daily (0 or 20 mg/kg, i.p.) for 12 consecutive days, and behavior recorded on days 1, 6, and 12. After 28 days without drug, rats were challenged with cocaine (20 mg/kg) and behavior was assessed. Novel reactivity phenotype had a dramatic role in adolescents’ acute responsiveness to cocaine. HR adolescents exhibited a robust increase in sensitivity to acute administration of 20 mg/kg cocaine. However, following repeated cocaine exposure, both LR and HR adults displayed a marked increase in cocaine-induced locomotor activity, whereas the change in behavioral responsiveness in LR and HR adolescents was not as dramatic. During the cocaine challenge trial, LR that were exposed to 20 mg/kg cocaine during adolescence showed the highest cocaine-induced responsiveness compared to their saline-exposed and adult-exposed counterparts. Overall, these data indicate that novel reactivity phenotype and age of initial cocaine exposure are intricately involved in the long-term behavioral effects of cocaine. Thus, novel reactivity phenotype may predispose cocaine-using adolescents to continue drug-seeking behavior and potential abuse.

**Highlights:**

- Novel reactivity is higher in adolescents compared to adults
- Adolescents show greater acute cocaine-induced activity compared to adults
- High responding adolescents show robust acute cocaine-induced activity
- Low responding adolescents show lasting changes in cocaine activity in adulthood

## 1.0 INTRODUCTION

Adolescence is characterized by a heightened vulnerability to engage in risk-taking behaviors including unprotected sexual intercourse, reckless driving, smoking cigarettes, and abusing drugs and alcohol (Eaton et al., 2008; Steinberg, 2008). Drug experimentation is most prevalent prior to adulthood, with early initiation predicting chronic use/abuse and dependence for some individuals (DeWit, Adlaf, Offord, & Ogborne, 2000). Yet, only 15-20% of overall drug users eventually develop an addiction (Anthony, Warner, & Kessler, 1994; Arenas et al., 2016), raising interest as to which individual differences factor into the potential for abuse. Identifying phenotypic characteristics that facilitate engagement in risky behaviors could predict patterns of drug use initiation and long-term addiction.

Risky and reckless behaviors occur more frequently during adolescence than during childhood and adulthood, with a peak increase in risk-taking behavior during mid-adolescence (Doremus-Fitzwater, Varlinskaya, & Spear, 2010; Somerville, Jones, & Casey, 2010; Steinberg, 2008; Zuckerman, Eysenck, & Eysenck, 1978). Enhanced risky behavior is attributed to the specific manner adolescents respond to salient environments and cues. Adolescents are hypersensitive to salient environmental cues and are more motivated to seek out rewards and novel experiences than young children and adults (Doremus-Fitzwater et al., 2010; Somerville et al., 2010). Thus, reckless and dangerous decisions are sometimes the consequence of adolescents seeking out pleasurable experiences.

The increase in reward-seeking during adolescence is likely mediated by several neurological events occurring in the developing brain during adolescence including synaptic pruning and increased innervations between prefrontal cortices and subcortical structures in the limbic system (Brenhouse, Sonntag, & Andersen, 2008; Paus, 2005; Steinberg, 2008). Prior to adulthood, an imbalance occurs between the cognitive control of the prefrontal cortex and the reward/salience of the limbic system, leading adolescents to make more emotionally driven decisions and to engage in more risky behaviors (Nelson, Leibenluft, McClure, & Pine, 2005; Somerville et al., 2010; Steinberg, 2008). As a result, adolescents perceive pleasurable experiences more positively than younger and older subjects (Arenas et al., 2016; Cauffman et al., 2010; Crews & Boettiger, 2009; Ernst et al., 2005; Geier & Luna, 2009; Steinberg, 2008). Together, these data suggest, adolescents are uniquely susceptible to risky behaviors.

Rodent models of risk-taking behaviors have been able to recapitulate this enhanced frequency of risky behaviors during adolescence (Douglas, Varlinskaya, & Spear, 2003; Stansfield & Kirstein, 2006; Stansfield, Philpot, & Kirstein, 2004). One construct used to describe risk-taking behaviors in rodents is defined as novelty-seeking and involves exploration of novel situations and objects (Arenas et al., 2016). Novel reactivity is a behavioral assay that identifies subjects as low or high responders according to locomotor activity [total distance moved (TDM)] in an unfamiliar environment, while novel object preference is a behavioral assay where subjects can approach or avoid an unfamiliar object. Adolescent rats demonstrate novel object preference (Douglas et al., 2003; Stansfield & Kirstein, 2006; Stansfield et al., 2004) and exhibit greater risk-taking behavior than adult rats (Macrì, Adriani, Chiarotti, & Laviola, 2002) by spending more time with a novel object and by exhibiting shorter latencies to approach the novel object. Novel reactivity has been extended to studies of psychomotor stimulants such as cocaine and amphetamine. Interestingly, the novel reactivity phenotype is predictive of an individual’s response to psychomotor stimulants, an effect independent of the route of administration (i.e., passive exposure or self-administration), and which may be associated with the rewarding effects of psychoactive drugs (Hooks, Colvin, Juncos, & Justice, 1992; Klebaur, Bevins, Segar, & Bardo, 2001). While there exists some variability in how novelty-seeking is measured (e.g., novel responding vs. novel object preference), dividing subjects into low responders (LR) and high responders (HR) has been useful for identifying endophenotypes susceptible to drug abuse (Arenas et al., 2016; Doremus-Fitzwater et al., 2010; Hittner & Swickert, 2006; Kelly et al., 2006; Stansfield et al., 2004). For instance, HR exhibit greater locomotor activation and self-administration in response to acute amphetamine and cocaine as compared to LR (Hooks, Jones, Smith, Neill, & Justice, 1991). In addition, HR are more sensitive to stress- and drug-primed reinstatement than LR (Bevins, Klebaur, & Bardo, 1997; Klebaur et al., 2001) and demonstrate a greater escalation of cocaine self-administration (Mantsch, Ho, Schlussman, & Kreek, 2001). While the lasting effects of chronic cocaine exposure have been observed in HR adults, the role of novel reactivity phenotype in determining cocaine responsiveness for adolescents remains uncertain. To better understand individual susceptibilities to drug abuse, novel reactivity and novel object preference were examined to determine endophenotypes that may be sensitive to repeated cocaine exposure during adolescence as compared to adulthood.

## 2.0 METHODS

### 2.1 Subjects

Adolescent (n=39) and adult (n=36) male Sprague Dawley rats were used for the present experiment. All rats were obtained from an in-house breeding colony at the University of South Florida, Tampa. Rats were sexed and culled to 10 pups on postnatal day (PND) 1. Pups remained with their respective dams until weaning. Only one male pup per litter was placed in any given treatment dose condition, and no more than two animals per litter were placed in the same treatment condition to later be separated based on novel reactivity phenotype. On PND 21 pups were weaned and pair-housed with same-sex littermates. Rats were maintained on a 12:12 hour light:dark cycle in a temperature and humidity-controlled vivarium. All rats were allowed free access to food and water throughout the experiment. Maintenance and treatment of all rats were within the guidelines for animal care by the National Institutes of Health (Public Health Service Policy on Humane Care and Use of Laboratory Animals, NIH 2002).

Animals at each age were separated into LR and HR utilizing a median split in which all animals were ranked according to total distance moved (TDM) on Trial 1 of the novel object preference paradigm (see below). Rats below the median were classified as LR and rats above the median were classified as HR. Utilizing this median split design, there was not an equal number of animals in each condition given that treatment dose was randomly assigned. Adolescent animals began on PND 31 and adults on PND 61. During the course of the experiment, adolescents and adults were tested between PND 31-74 and PND 61-104, respectively.

### 2.2 Novel object preference paradigm

A schematic of the experimental procedure can be seen in Figure 1. To assess novel reactivity phenotype, rats (PND 31-34 for adolescents and PND 61-64 for adults) were placed in an open arena to measure spontaneous activity levels in a novel environment. The open arena was composed of a black plastic circular platform with a white plastic circular barrier that enclosed the testing area (diameter = 96 cm). A white plastic curtain further enclosed the testing area to minimize spatial cues. Movement was tracked using a camera suspended above the open arena and linked to the Ethovision Behavioral Tracking System (Noldus Information Technology, Utrecht, the Netherlands). On the morning of the 1st trial, all rats were introduced into the novel open field for 5 min. Total distance moved (TDM, cm) during this trial was used to identify novel reactivity phenotype as low or high levels of spontaneous activity in a novel environment (novel reactivity). Trials continued twice daily for 4 days to habituate rats to the environment. After the 8th trial on the afternoon of the 4th day, rats were removed from the open arena for one min, the apparatus was cleaned and a novel object (approximately 16.5 cm tall) was placed in the center of the table using a magnet to keep it in place. During the 9th trial the animal was placed back into the open arena for 5 min and time spent in proximity to the novel object, frequency to approach the novel object, latency to approach the novel object, and TDM were measured (novel object preference).

**Figure 1:**
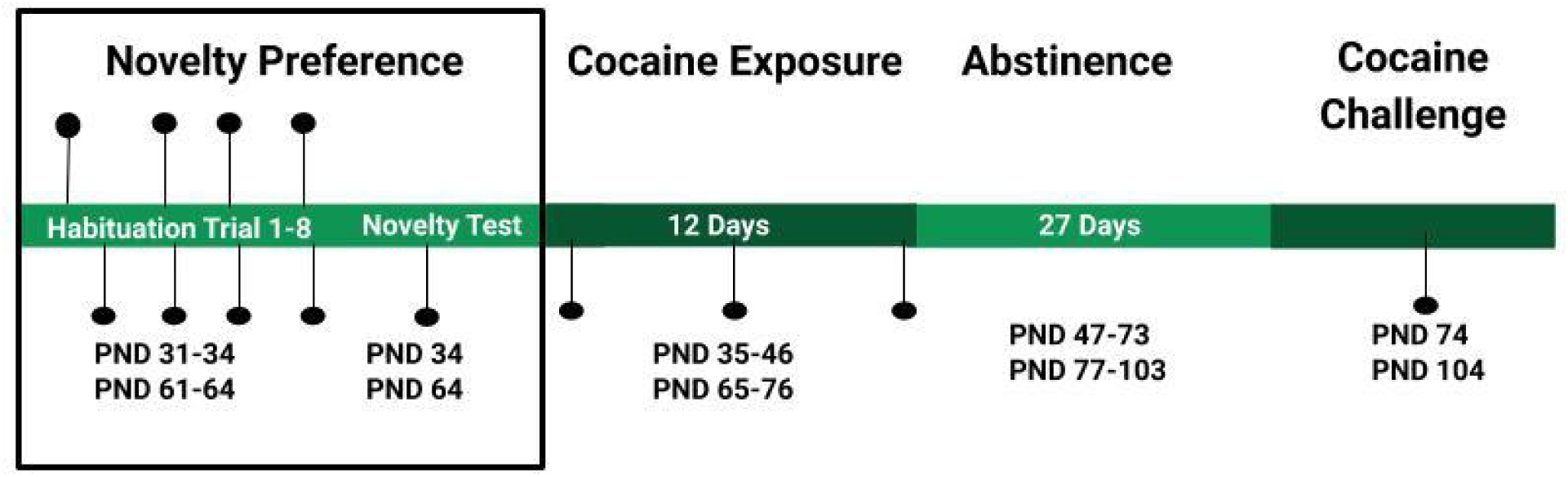
Experimental Timeline. Animals underwent the novel object preference habituation trials and test from PND 31-34 for adolescents and PND 61-64 for adults. This was followed by 12 days of cocaine or saline exposure from PND 35-46 for adolescents and from PND 65-76 for adults. All rats were tested for cocaine-induced TDM on PND 35, 40, and 46 for adolescent-exposed animals and on PND 65, 70, and 76 for adult-exposed animals. On the 28th day after the last cocaine injection, all animals were administered 20 mg/kg of cocaine on PND 74 or PND 104 for adolescent-exposed rats and adult-exposed rats, respectively.

### 2.3 Cocaine Exposure

After completion of the novel object preference test, rats were injected daily with cocaine (20 mg/kg) or vehicle (0.9% NaCl) for 12 consecutive days. Adolescents were injected on PND 35-46 and adults were injected on PND 65-76. Doses were administered at a dose of 1 ml/kg.

#### 2.3.1 Cocaine-induced Total Distance Moved Trials

During the 12-day cocaine exposure, locomotor behavior was assessed across trials as total distance moved (TDM) for adolescents (PND 35, 40 and 46) and adults (PND 65, 70 and 76). For each TDM trial, rats were placed in an open arena for 30 min and left to explore the entire arena. After the first 30 min of the trial, rats were removed from the arena and received their respective injection of 20 mg/kg cocaine or vehicle and immediately placed back in the arena for an additional 45 min. While in the arena, movement was constantly tracked and quantified using Noldus software.

#### 2.3.2 Cocaine Challenge

After cocaine exposure, rats entered a 27-day (PND 47-73 or PND 75-103) abstinence period in which rats were not manipulated except for regular cage changes. After the abstinence period, on PND 74 or PND 104, all rats were challenged with 20 mg/kg cocaine and TDM was assessed. Behavior tracking was identical to that described above in novel object preference paradigm.

### 2.4 Design and Analyses

To assess differences in novel reactivity phenotype, data for TDM across the eight habituation trials and during the novel object preference test (trial 9) were analyzed using a two factor ANOVA with Age (2; Adolescent, Adult) as a between subjects factor and trial (9) as a repeated measure. For the novel object preference test, data for time spent with, frequency to approach, and latency to approach the novel object were analyzed utilizing a two-factor between subjects design ANOVA for Age (2; Adolescent, Adults) and Responder (2; LR, HR).

During the cocaine exposure behavioral assessment trials, habituation and cocaine-induced TDM data were analyzed separately using a four-factor mixed model ANOVA with Age (2; Adolescent, Adult), Dose (2; 0, 20) Responder (2; LR, HR) as between subjects factors and Trial (3; Trial 1, Trial 6, Trial 12) as a repeated measure. Data for the cocaine Challenge trial were analyzed using a 3 factor between subjects design ANOVA for Age (2: Adolescent, Adult), Dose (2; 0, 20), and Responder (2: LR, HR).

Time course data for each trial were analyzed separately for LR and HR using a mixed-model design ANOVA for Age (2: Adolescent, Adult) and Dose (2; 0, 20) as between subjects factors with Time (9; 5, 10, 15, 20, 25, 30, 35, 40, 45 min) as a repeated measure. In the event of significant interactions, post hoc analyses utilized Tukey HSD tests to isolate effects. The level of significance was set at α ≤ 05.

## 3.0 RESULTS

### 3.1 Novel Environment Induced TDM/Preference Test

#### 3.1.1 Novel Object Preference Habituation Trials

Rats were separated into LR and HR utilizing a median split based on TDM on the first 5 min trial in the novel open field. Rats were ranked according to TDM on trial 1 within age, and regardless of dose, separated into their respective responder groups. Given that exposure dose was randomly assigned, there was an unequal number of animals in each condition (see Fig. 2 legend).

**Figure 2:**
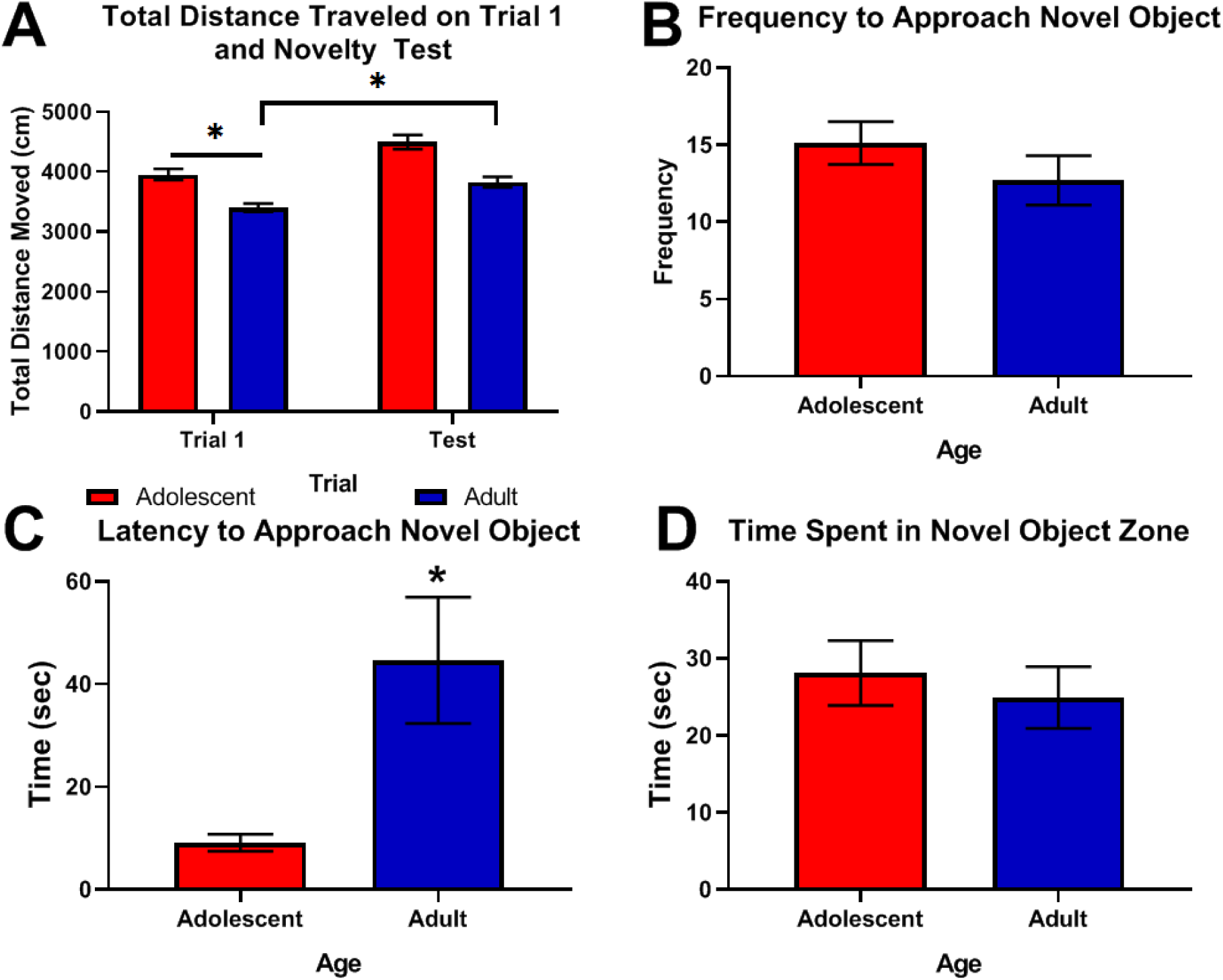
Data depict mean +/− SEM for (Panel A) total distance moved (cm) during each habituation trial when animals were exposed to the open field and (Panel B) frequency to approach the novel object, (Panel C) time spent in the novel zone (sec), and (Panel D) latency to approach the novel object (sec). Adolescent LR n=20, Adolescent HR n= 19, Adult LR n=16, Adult HR n=20. * indicates Adolescent is significantly greater than Adult. ^ indicates TDM for LR is greater on the test trial than on habituation trials 1 and 2. ^^ indicates HR is significantly greater than LR.

As depicted in Figure 2A, adolescent rats exhibited greater variability in locomotor activity across the eight habituation trials and during the novel object preference test on trial 9, compared to adults as supported by an Age X Trial interaction [*F*(8, 528) = 5.94, p<0001] and main effects of Age [*F*(1, 66) = 26.22, p<0005] and Trial [*F*(8, 528) = 4.40, p<0001]. Specifically, adolescent rats showed higher TDM compared to adults on trials 4-6, trial 8, and during the novel object preference test on trial 9 (all p < 001). Additionally, LR rats showed higher TDM on the test trial compared to the first two habituation trials (all p< 05) as supported by a Trial X Responder interaction [*F*(8, 528) = 3.33, p<01] and main effect of Responder [*F*(1, 66) = 8.45, p<005].

#### 3.1.2 Novel Object Preference Test Trial

Using the same separation for designation in LR and HR based on TDM for trial 1, assessing animals for frequency to approach (Figure 2B), HR rats approached the novel object more often compared to LR as supported by a main effect of Responder [*F*(1, 67) = 7.27, p<01]. When measuring time spent with the novel object (Figure 2C) and latency to approach the novel object (Figure 2D), there were no statistically significant differences between LR and HR or between adolescents and adults.

### 3.2 Cocaine-Induced Locomotor Activity

#### 3.2.1 Habituation

When data were analyzed for Age, Dose, and Responder across the Exposure and Cocaine Challenge trials, there were no differences in rate of habituation to the environment (data not shown). However, TDM during the habituation period was lowest during the cocaine challenge trial compared to all other trials as supported by a main effect of Trial [*F*(1, 130) = 9.51, p < 0001].

#### 3.2.2 Cocaine-induced TDM

When data were analyzed for TDM across the cocaine exposure trials, there were Age, Dose, and Responder differences. Specifically, adolescent rats administered cocaine showed higher cocaine-induced TDM compared to adults administered cocaine during Trial 1 (Figure 1A), but not during Trial 6 (Figure 1B) or Trial 12 (Figure 1C), as supported by an Age X Dose X Trial [*F*(2, 130) = 4.39, p < 02] interaction. In adolescents, there were no differences in cocaine-induced TDM across trials, but the adults showed higher TDM at Trial 6 (Figure 1B) and Trial 12 (Figure 1C) compared to Trial 1.

Post hoc analyses of the Responder X Trial interaction [*F*(2, 130) = 3.08, p < 05] did not reveal any significant differences. The Age X Trial [*F*(2, 130) = 5.31, p < 01] interaction showed that adolescents had higher TDM compared to adults on Trial 1, but not Trials 6 or 12. Adolescents administered cocaine had higher TDM compared to cocaine administered adults as supported by an Age X Dose [*F*(1, 65) = 11.21, p < 005] interaction and main effects of Age [*F*(1, 65) = 5.65, p < 05] and Dose [*F*(1, 65) = 225.32, p < 001].

#### 3.2.3 Cocaine Challenge Test

During the challenge administration of cocaine test (Figure 2), there were effects of age and cocaine exposure on cocaine-induced TDM. Specifically, rats that were exposed to repeated cocaine and challenged with 20 mg/kg of cocaine showed higher TDM compared to animals that were exposed to saline and administered acute 20 mg/kg cocaine for the first time at the Challenge test, as supported by a main effect of Dose [*F*(1, 67) = 20.74, p < 0005]. Rats that were exposed during adolescence showed higher TDM during the cocaine challenge test compared to those that were exposed during adulthood, as supported by a main effect of Age [*F*(1, 67) = 5.4, p < 05].

### 3.3 Time Course Analyses

#### 3.3.1 Low Responders

To determine the temporal phenotypic response of LR across exposure, data were analyzed at each trial for Age X Dose X Time. For LR, at the first cocaine exposure during Trial 1 (Figure 3A), adolescents showed higher cocaine-induced TDM compared to adults as supported by a significant Age X Dose interaction [*F*(1, 32) = 4.31, p < 05] and a main effect of Dose [*F*(1, 32) = 47.81, p < 0001]. All animals administered cocaine showed higher cocaine-induced TDM compared to saline-administered animals at every time point supported by a Time X Dose interaction [*F*(8, 256) = 4.39, p<0001] and a main effect of time [*F*(8, 256) = 12.32, p<0001].

**Figure 3:**
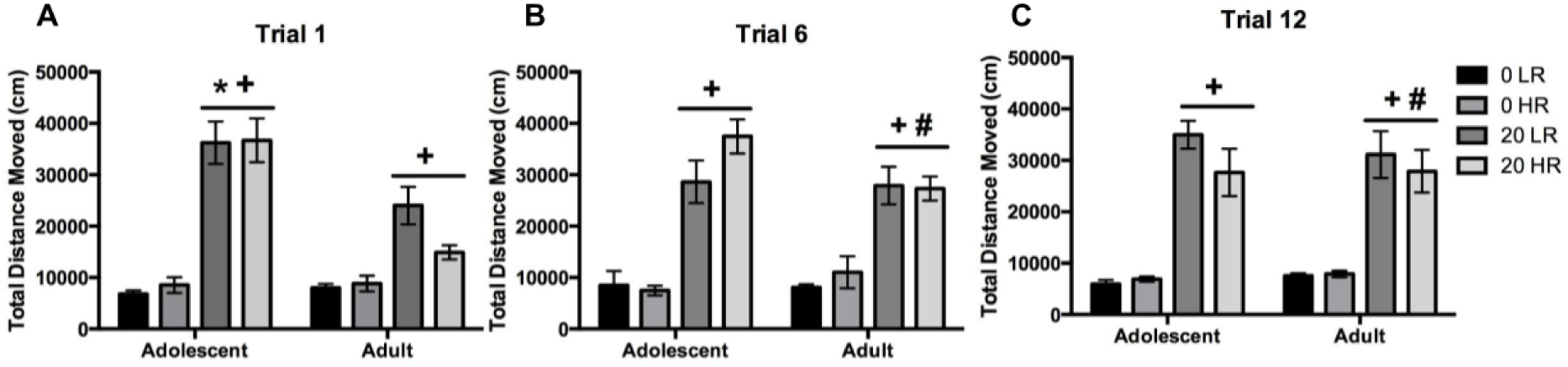
Data depict mean +/− SEM for total distance moved (cm) for the entire 45 min trial during Trial 1 (Panel A), Trial 6 (Panel B), and Trial 12 (Panel C) for animals administered saline or 20 mg/kg cocaine. Adolescent LR 0 n=8, Adolescent LR 20 n=12, Adult LR 0 n=8, Adult LR 20 n=8, Adolescent HR 0 n=11, Adolescent HR 20 n=8, Adult HR 0 n=10, Adult HR 20 n=10. * indicates Adolescent is significantly greater than Adult. + indicates cocaine is greater than saline. # indicates TDM adults administered cocaine is significantly greater on Trials 6 and 12 compared to Trial 1.

During trial 6 (Figure 3B) and Trial 12 (Figure 3C), LR that were administered cocaine showed increased TDM across time as supported by main effects of Dose [[Trial 6: [*F(*1, 32) = 41.57, p < 001]; Trial 12: [*F*(1, 32) =89.42, p < 0001] and Time [Trial 6: [*F*(8, 256) = 5.33, p < 0001]; Trial 12: [*F*(8, 256) = 7.84, p < 0001]].

During the Cocaine Challenge test (Figure 3D), LR that were repeatedly exposed to cocaine showed higher TDM compared to LR that were repeatedly exposed to saline at 25 min (p = 054), 30 min (p< 05), and 40 min (p< 02) of the trial as supported by a Dose X Time interaction [*F*(8, 256) = 2.10, p < 05]. Additionally, main effects of Age [*F*(1, 32) = 6.97, p < 05,], Dose [*F*(1, 32)= 10.69, p < 005], and Time [*F*(8, 256) = 18.71, p < 0001] suggest that animals exposed to cocaine during adolescence and challenged with cocaine as adults showed increased TDM compared to similarly exposed adults and animals that were exposed to saline during adolescence.

#### 3.3.2 High Responders

During the initial exposure to cocaine during Trial 1 (Figure 5A), TDM varied as a function of Age, Dose, and Time. Specifically, adolescent HR that were exposed to cocaine showed higher TDM compared to adult HR that were exposed to cocaine from 5 min to 30 min of the trial (all p<01) as supported by Age X Dose X Time [*F*(8, 272) = 5.00, p<0001], Age X Time [*F*(8, 272) = 5.71, p<001], and Dose X Time interactions [*F*(8, 272) = 3.60, p<001] and main effect of Time [*F*(8, 272) = 25.54, p<0001]. Additionally, adolescents HR rats showed the highest increase in TDM during the acute cocaine exposure compared to all other groups as supported by Age X Dose interaction [*F*(1, 34) = 24.58, p<0001] and main effects of Age [*F*(1, 34) = 23.57, p<0001], and Dose [*F*(1, 34) = 59.27, p<0001].

During Trial 6 (Figure 4B), all animals administered cocaine had greater TDM compared to their saline-exposed counterparts, and this effect was the highest in the cocaine-exposed adolescent HR supported by a significant Age X Dose [*F*(1, 34) = 4.42, p<05], Time X Dose interaction [*F*(8, 272) = 2.33, p<05] and main effects of Dose [*F*(1, 34) = 63.99, p<0001] and Time [*F*(8, 272) = 11.19, p<0001]. During Trial 12 (Figure 4C), adolescent HR administered cocaine showed the higher cocaine-induced TDM compared to their saline-exposed counterparts at 5 min, and from 25-40 min of the trial, and adult HR had higher cocaine-induced TDM compare to saline-exposed controls from 5-20 min and at 45 min of the trial (all p < 05) as supported by a Age X Dose X Time [*F*(8, 280) = 3.14, p<005], Age X Time [*F*(8, 280) = 4.89, p<0001], Dose X Time [*F*(8, 280) = 4.95, p<0001] interactions and main effects of Dose [*F*(1, 35) = 49.58, p<001] and Time [*F*(8, 280) = 20.39, p<0001]. There were no age of exposure differences between adolescents and adults administered saline nor between adolescents and adults administered cocaine during Trial 12. During the Challenge test (Figure 4D), all HR animals administered cocaine showed higher TDM across time compared to saline-administered controls as supported by a main effects of Dose [*F*(1, 35) = 10.33, p<005] and Time [*F*(8, 280) = 23.77, p<001].

**Figure 4:**
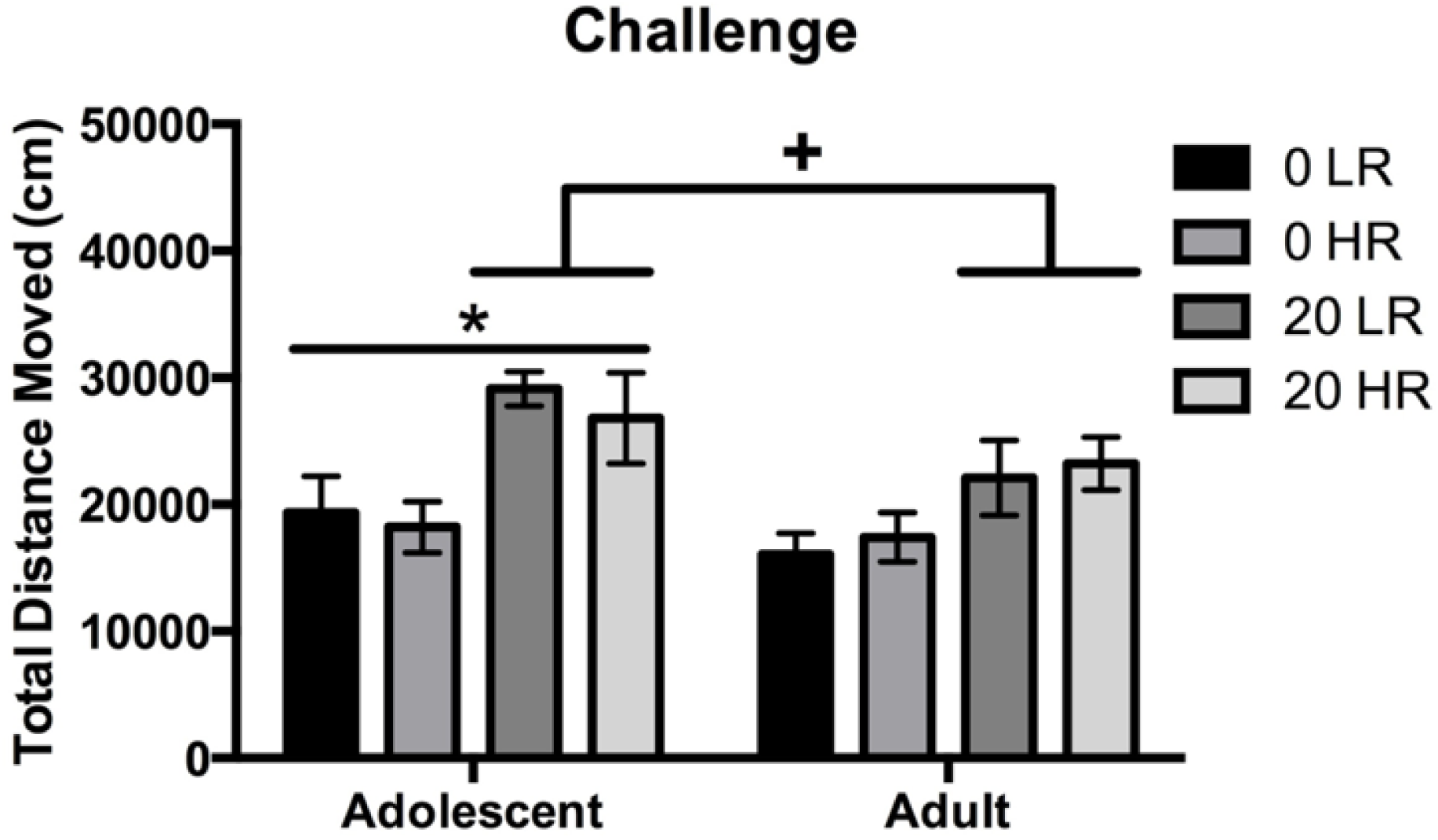
Data depict mean +/− SEM for total distance moved (cm) in response to 20 mg/kg cocaine for the entire 45 min trial during Cocaine Challenge trial. * indicates Adolescent is significantly greater than Adult. + indicates cocaine is greater than saline.

## 4.0 DISCUSSION

### 4.1 Overview

The primary focus of the present study investigated the influence of novel reactivity phenotype on the long-term behavioral effects of cocaine exposure in adolescents compared to adult male rats. To our knowledge, this is the first experiment to investigate the influence of innate novel reactivity phenotype in adolescent and adult male rats on immediate and long-term changes in cocaine responsiveness. The present experiment replicates and extends the literature indicating adolescents respond to the acute effects of cocaine with heightened locomotor behavior (Badanich, Maldonado, & Kirstein, 2008; Catlow & Kirstein, 2005; A. Maldonado & C. L. Kirstein, 2005; A. M. Maldonado & C. L. Kirstein, 2005; McQuown, Dao, Belluzzi, & Leslie, 2009; Schramm-Sapyta, Pratt, & Winder, 2004), while adults gradually increase their locomotor behavior over time in response to repeated cocaine administration (Collins & Izenwasser, 2002; Frantz, O’Dell, & Parsons, 2007). Only LR adolescent animals exposed and challenged with 20 mg/kg cocaine maintained greater cocaine-induced locomotor behavior relative to LR adults exposed and challenged with the same dose. The cocaine challenge data reveal that novel reactivity phenotype and age of initial cocaine exposure are intricately involved in the long-term behavioral effects of cocaine.

### 4.2 Novel object preference phenotype

In previous work, novel reactivity was the most reliable predictor for long-term response to repeated methylphenidate in adolescent and adult rats compared to free-choice novelty preference (Wooters, Dwoskin, & Bardo, 2006). The present experiment utilized an overall median split design based on TDM in an inescapable novel environment. Similar to other work, HR adolescents had the highest level of novel environment-induced reactivity relative to all other groups, followed by HR adults and LR adolescents when first exposed to a novel environment (Philpot & Wecker, 2008; Stansfield & Kirstein, 2005; Stansfield et al., 2004).

It is important to note that the LR/HR separation parameters used in this study, relative to different novel object preference separation parameters (i.e., latency to approach novel object, frequency to approach novel object, and time spent with novel object), could lead to different results (Bardo, Donohew, & Harrington, 1996; Wooters et al., 2006). For example, similar to the work presented here, Wooters and colleagues found that a free-choice novelty test did not predict the effect of methylphenidate, while inescapable novel environment-induced TDM did predict the effects of repeated drug administration for saline and drug challenge days (Wooters et al., 2006). There were no significant correlations between the novel object preference parameters and cocaine-induced TDM on Trial 1 or during the cocaine Challenge test (data not shown).

### 4.3 Cocaine exposure

Novel reactivity phenotype influenced the acute response to 20 mg/kg of cocaine in adolescents. Specifically, HR adolescents showed robust increases in cocaine-induced TDM across time as compared to HR adults (Figure 6). In contrast, LR adolescents showed a similar profile, but the increases in cocaine-induced TDM was less robust across time as compared to LR adults (Figure 5). In addition, other work supports our findings that cocaine-induced TDM is greater for adolescent than adult rats (Badanich et al., 2008; Catlow & Kirstein, 2005; McQuown et al., 2009). In contrast, other studies suggest that adolescents show hyporeactivity to psychostimulant drug-induced TDM compared to adults (Collins & Izenwasser, 2002; Laviola, Adriani, Terranova, & Gerra, 1999; Spear, 2000; Torres-Reveron & Dow-Edwards, 2005; Wiley, Evans, Grainger, & Nicholson, 2008; Wooters et al., 2006), which was not observed in the present experiment.

**Figure 5:**
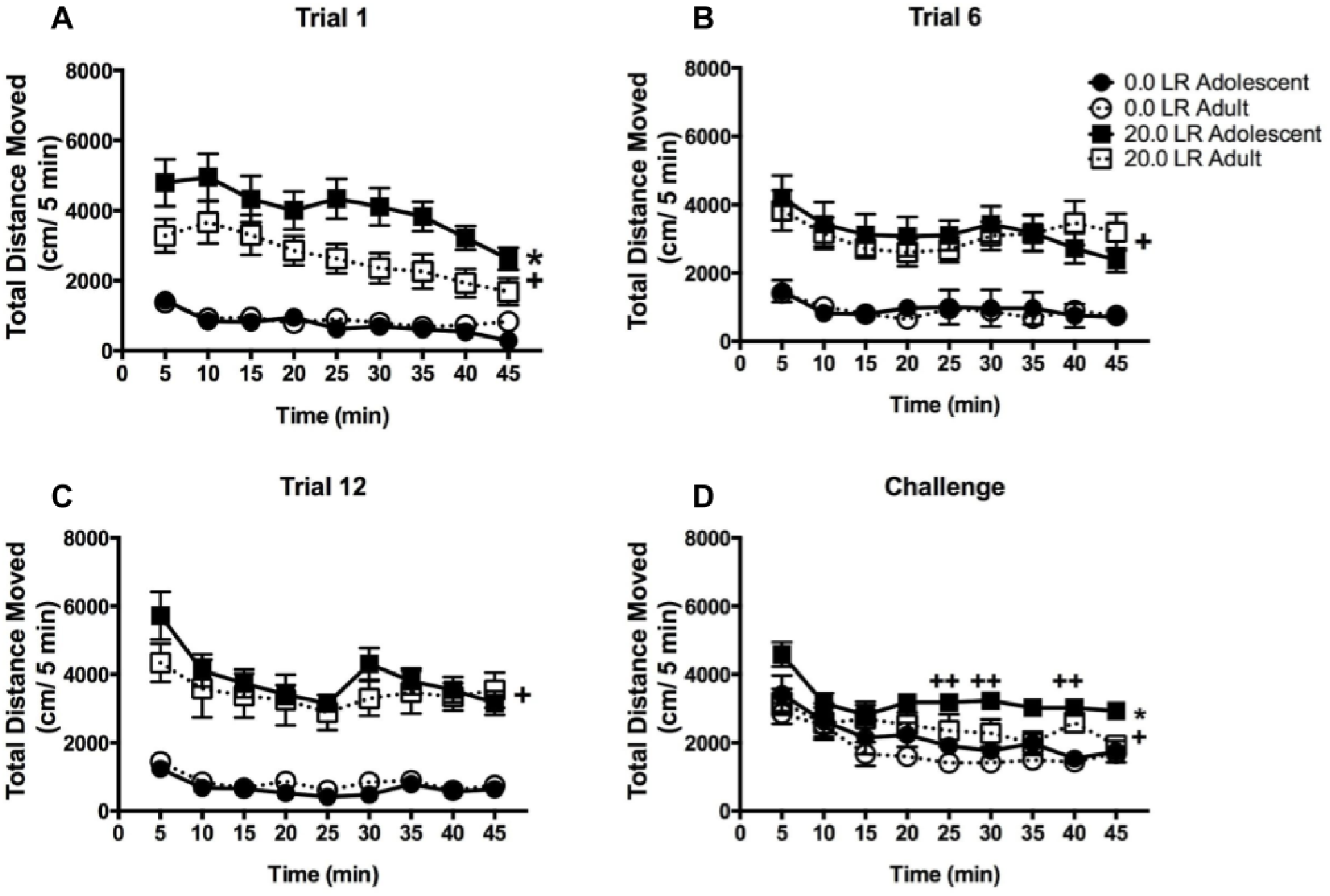
Data depict mean +/− SEM for total distance moved (cm) for LR during the post-injection period in 5 min intervals on (Panel A) Trial 1, (Panel B) Trial 6, (Panel C) Trial 12 of saline or cocaine exposure and (Panel D) following cocaine challenge administration in adolescent and adult rats. * indicates Adolescent is significantly greater than Adult. + indicates cocaine is greater than saline. ++ indicates adolescent administered cocaine is significantly greater than adolescent administered saline.

**Figure 6:**
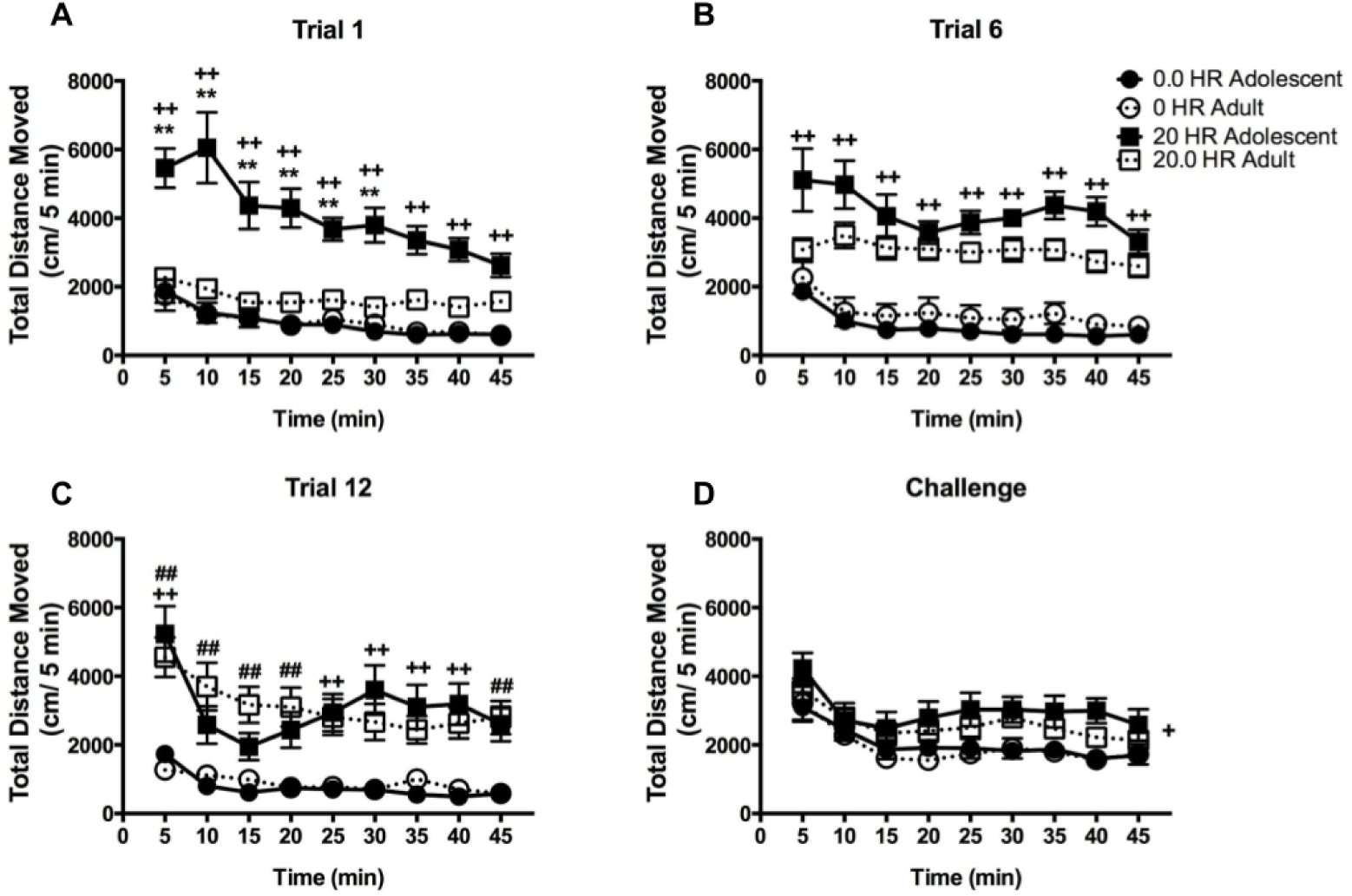
Data depict mean +/− SEM for total distance moved (cm) for HR during the post-injection period in 5 min intervals on (Panel A) Trial 1, (Panel B) Trial 6, (Panel C) Trial 12 of saline or cocaine exposure and (Panel D) following cocaine challenge administration in adolescent and adult rats. ** indicates adolescents administered cocaine is significantly greater than adults administered cocaine ++ indicates adolescent HR administered cocaine is significantly greater than adolescent HR administered saline. ## indicates adults administered cocaine is significantly greater than adults administered saline. + indicates cocaine is greater than saline.

When comparing age, both LR adolescents and LR adults showed similar cocaine-induced TDM by the 6th administration of cocaine. In contrast, HR adolescents administered 20 mg/kg of cocaine did not show similar cocaine-induced TDM relative to HR adults until Trial 12. When comparing within age, the adults exposed to 20 mg/kg of cocaine showed heightened drug-induced TDM on Trial 6 and 12 relative to the first day of treatment. Others support this observation that adults repeatedly exposed to cocaine show greater TDM over 7 to 15 days of cocaine exposure (Collins & Izenwasser, 2002; Frantz et al., 2007). Together, these findings indicate the behavioral responsiveness to repeated cocaine is different for adolescent and adult rats, an effect which is modulated by novel reactivity phenotype.

### 4.4 Challenge

Data showing the long-term effects of psychostimulant administration during adolescence on the responsivity to a drug challenge in adulthood vary. Similar to the present work, adolescent mice exposed to 10 mg/kg of cocaine displayed elevated TDM, relative to adults, in response to a cocaine challenge administration of the same dose (Camarini, Griffin, Yanke, dos Santos, & Olive, 2008). Marin and colleagues (Marin, Cruz, & Planeta, 2008) showed that 3 and 30 days after cocaine exposure (10 mg/kg b.i.d. from PND 30-34), there was a significant increase in cocaine-induced activity in adolescent exposed rats. This effect was not observed after 60 days without cocaine, which is indicative of lasting effects of cocaine pretreatment during adolescence. In addition, adolescent rats with a history of oral cocaine consumption showed decreased cocaine-induced TDM to a challenge administration of cocaine during adolescence (Walker et al., 2009). Previous work has shown that chronic exposure to methylphenidate during adolescence predisposes rats to exhibit greater responsiveness to cocaine self-administration after a two-week abstinence period (Brandon, Marinelli, Baker, & White, 2001). Nicotine pretreatment during adolescence, but not adulthood, sensitized rats to cocaine-induced elevations in TDM when challenged three days later, an effect that was likely potentiated by novelty (McQuown et al., 2009). Thus, data regarding the long-term effects of psychostimulant administration on reactivity to a drug challenge in adulthood are mixed, and are likely mediated by a multitude of factors (e.g., age of exposure, dose of pretreatment, dose during drug challenge, and time between initial pretreatment and challenge).

It is also possible that the challenge dose of cocaine used in the present experiment induced a ceiling effect in cocaine-induced TDM and if a lower dose of cocaine had been used during the cocaine challenge test, differences in behavior may have been observed as was shown in preweanling rats pretreated with 30 mg/kg and challenged with 15 mg/kg cocaine in a context-dependent treatment regimen (Wood et al., 1998). Moreover, the abstinence period between cocaine treatment and challenge cocaine administration may have been too long to observe any lasting changes following cocaine pretreatment in the adolescent rats (Zavala, Nazarian, Crawford, & McDougall, 2000). Nonetheless, despite these potential limitations, the present work demonstrates a unique interaction between innate novel reactivity phenotype and long-term responsiveness to a high dose of cocaine in adolescents compared to adult rats.

### 4.5 Conclusion

The present data suggest that adolescent male rats are more vulnerable to the effects of repeated cocaine exposure than adult male rats. Acute cocaine-induced activity in response to 20 mg/kg cocaine was similar in LR and HR adolescent rats, followed by elevated activity in LR adult rats and the lowest cocaine-induced responsiveness in HR adult rats. These data indicate novel reactivity phenotype plays a dramatic role in overall acute adult cocaine-induced TDM and less of a role in overall acute adolescent cocaine-induced TDM. Adolescents exposed to 20 mg/kg of cocaine developed a unique bimodal curve in cocaine-induced locomotor behavior, an effect that was absent in adult animals. The cocaine challenge data suggest that pre-exposure to 20 mg/kg cocaine induced lasting sensitivity to subsequent cocaine challenge. Results from the cocaine challenge trial indicate LR adolescents are the most sensitive to the lasting changes induced by drug exposure during adolescence, as supported by the overall high level of TDM compared to LR adults exposed to 20 mg/kg cocaine. These data support the emerging hypothesis that LR animals are more sensitive to the long-term effects of cocaine. If these data were to be extrapolated to humans, the LR or low sensation-seeking human would be less likely to initiate use of cocaine, and thus this phenotype would serve as a protective mechanism against initial cocaine use. However, if the LR adolescents were to begin using cocaine, they could develop similar behavioral responsiveness to the long-term consequences induced by cocaine. Taken together, these data indicate that adolescent male rats are both behaviorally and neurologically more vulnerable to the effects of cocaine than adult male rats. This study adds to the literature, demonstrating that innate novel reactivity phenotype has different long-term implications that alter responsivity to repeated cocaine during adolescence or adulthood. Additional studies in both males and females should be conducted across age to further understand the neurobiological basis of novel reactivity phenotype and behavioral responsivity to both short-term and long-term use of cocaine.

## Acknowledgements

This work was supported by the University of South Florida Department of Psychology and the College of Arts and Sciences.

